# Cerebrovascular response dynamics to hypercapnia in healthy aging

**DOI:** 10.1101/2025.10.01.679768

**Authors:** Mauro DiNuzzo, Maria Guidi, Giovanni Giulietti, Edoardo D’Andrea, Taljinder Singh, Matteo Mancini, Claudia Marzi, Marco Clemenzi, Laura Serra, Federico Giove

**Affiliations:** Enrico Fermi Research Center, Rome, Italy; Neuroimaging Laboratory, Fondazione Santa Lucia IRCCS, Rome, Italy; National Institute for Nuclear Physics – National Laboratories of the South (INFN-LNS), Catania, Italy

**Keywords:** neurovascular coupling, cerebral blood flow, computational modeling, vascular biomarkers, functional neuroimaging

## Abstract

Cerebrovascular dysfunction is an early and underrecognized contributor to cognitive decline. Standard measures such as cerebrovascular reactivity (CVR) during hypercapnia capture only the amplitude of flow responses, providing limited insight into the timing of vascular adaptation. Temporal features, such as delay (onset latency) and time constant (rate of adjustment), together with gain (response amplitude) may serve as more sensitive indicators of vascular health, but cannot be directly obtained from conventional imaging. Here, we investigated cerebral blood flow (CBF), cerebral blood volume (CBV), and blood oxygenation level dependent (BOLD) signal dynamics during hypercapnic challenge in healthy aging. Using a physiologically validated computational model, we estimated delay, time constant, and gain by optimizing the mapping of end-tidal gases to their arterial counterparts in a region-of-interest framework. Once parametrized using CBF, the model successfully predicted CBV and BOLD responses in independent experimental sessions. Across subjects, aging was associated with widespread heterogeneous region-specific changes in delay and substantial reductions in gain and time constant, indicating that cerebrovascular responses become weaker and less adaptable with age. These results demonstrate that calibrated simulations have the ability to track vascular aging, allowing the extraction of parameters that may represent novel biomarkers of cerebrovascular dysfunction. Unlike conventional CVR, temporal hemodynamic parameters capture the dynamics of vascular adaptation, providing a complementary dimension for early detection and therapeutic monitoring in aging and disease.

## Introduction

Cerebral blood flow (CBF) is tightly regulated to match the energetic and homeostatic demands of neuronal activity within a process termed neurovascular coupling that underlies the contrast exploited by functional neuroimaging (Burma et al., 2025; Tournissac et al., 2024). Disturbances in vascular regulation are increasingly implicated as contributors to aging, and cerebrovascular dysfunction can precede, interact with, and accelerate cognitive decline and dementia (Zlokovic et al., 2020). Importantly, alterations in the temporal profile of vascular responses may emerge before gross structural or metabolic loss, suggesting that hemodynamic metrics could provide sensitive, early indicators of cerebrovascular impairment (Marchena-Romero et al., 2023; Nanayakkara et al., 2025). Magnetic resonance imaging (MRI) methods are commonly used to probe cerebrovascular reactivity (CVR) in combination with vasoactive challenges, such as hypercapnia or breath-holding. While CVR has become a widely used metric of vascular health, it largely reflects the magnitude of flow responsiveness. By contrast, the temporal features of the response, including the delay (latency to onset) and the time constant (rate of adjustment to a new steady state), remain poorly characterized in vivo (Poulin et al., 1997; Poulin et al., 1996), yet they may carry critical information to probe cerebrovascular function (Turner et al., 2022).

Computational modeling has emerged as a powerful strategy to interpret and extend experimental measures of brain physiology (Aubert & Costalat, 2002; Aubert et al., 2007; Blanchard et al., 2016; Cloutier et al., 2009; DiNuzzo et al., 2011). Existing models of the BOLD response and vascular regulation have provided valuable insights, but most treat vascular dynamics with simplified assumptions, limiting their ability to capture fine-grained hemodynamic features from experimental data. On the other hand, experimental quantification of temporal vascular parameters remains challenging, as available imaging methods often lack the required temporal resolution or physiological specificity (Gupta et al., 2012; Ottoy et al., 2025).

In addition to nutrients supply, recent advancements in neurovascular research have highlighted the intricate relationship between cerebral blood flow (CBF) dynamics and metabolic waste clearance in the brain (Holstein-Rønsbo et al., 2023; Im et al., 2025; Rowsthorn et al., 2025; Wang et al., 2022). Structural alterations in the cerebral vasculature, including vessel varicosities and other forms of vascular remodeling, have indeed been implicated in impaired waste clearance mechanisms (Krings et al., 2025; Mestre et al., 2017). By perturbing CBF responses, these structural changes can potentially compromise the timely delivery of nutrients as well as the efficient removal of metabolic byproducts like CO_2_ and protons, thereby contributing to neuronal stress and functional decline across aging (DiNuzzo et al., 2024; Mangia et al., 2025). Understanding whether and how hemodynamic parameters change during healthy aging is particularly important. Cerebrovascular dysfunction has been implicated in aging as well as in the onset and progression of cognitive decline, with evidence for impaired vasoreactivity, endothelial (i.e., blood brain barrier) dysfunction, and reduced neurovascular coupling efficiency (Cantin et al., 2011; Iturria-Medina et al., 2016; Kisler et al., 2017; Kurz et al., 2022; Nehra et al., 2022; Nielsen et al., 2017; Sweeney et al., 2018). It is therefore reasonable to hypothesize that the dynamics of CBF regulation, not just its amplitude, degrade across the aging process.

In this study, we address this question by applying a biophysically detailed computational model of neurovascular and neurometabolic coupling to estimate delay, time constant, and gain of the brain vasculature based on the CBF response to hypercapnia in healthy aging subjects. The ability of the model to reproduce canonical cerebrovascular responses was validated against experimental changes in cerebral blood volume (CBV) and blood oxygenation-dependent level (BOLD) signal. By quantifying alterations in the temporal dynamics of vascular regulation across groups, we aim to provide novel insights into the vascular contributions to aging, and to evaluate the potential of hemodynamic parameters as sensitive biomarkers of vascular health.

## Materials and Methods

### Model development

To predict neuroimaging variables during neuronal activation and hypercapnic challenges, we have integrated several previously published mathematical and ordinary differential equation (ODE)-based models that have been used to simulate brain cellular metabolism and vascular regulation (see below). The integrated computational model consists of ten compartments (**Supplementary Table 1**) representing key physiological regions: arterial blood, precapillary arteriolar smooth muscle cells, capillary blood, capillary endothelium, endothelial basal lamina, interstitial space, glial cells, presynaptic neurons, postsynaptic neurons, and venous blood. The model includes approximately 80 molecular species (**Supplementary Table 2**) corresponding to the model balance equations forming the ODE system, 90 biochemical reactions and transport processes corresponding to the model rate equations (**Supplementary Table 3**), and about 150 parameters (**Supplementary Table 4**). Signaling mechanisms are implemented using a short-term plasticity (STP) model (DiNuzzo & Giove, 2012), incorporating sodium and potassium concentrations, compartment-specific Na+/K+ ATPase isoforms, and pH-dependent modulation of postsynaptic currents. Sodium and potassium currents are modeled using quantitative stoichiometry derived from a previous flux balance analysis (FBA) model (DiNuzzo et al., 2017). Cellular metabolism follows a framework adapted from previous models (Aubert et al., 2002; DiNuzzo et al., 2010), further incorporating explicit representation of protons, carbon dioxide, bicarbonate, and carbonic anhydrase-catalyzed reactions. Cerebral blood flow (CBF) regulation is modeled by adapting an existing framework (DiNuzzo et al., 2011; Yücel et al., 2009), accounting for direct neurogenic nitric oxide (NO)-mediated vasodilation and an indirect component via tissue CO_2_ diffusion to smooth muscle cells. Transport mechanisms are modeled using previous models (Aubert & Costalat, 2002; Simpson et al., 2007), further incorporating dissolved CO_2_ fluxes, anion exchange for the bicarbonate buffer system, and glial potassium buffering activity.

The model was parametrized for the human cerebral cortex and reproduces physiological values under resting conditions for known metabolite concentrations, conforming to previous validation works (e.g., Aubert et al., 2007). Moreover, the model qualitatively and quantitatively reproduces the outcomes of previously published models with respect to metabolite dynamics during stimulation (see Supplementary Information for details). The system of ODEs was numerically solved using the open-source Sundials CVODES solver (Lawrence Livermore National Laboratory) for stiff problems implemented in Matlab (The Mathworks, Inc.) version R2025a. The full model (balance equations, rate equations, and parameter values) is detailed in Supplementary Information ^1^.

### Experimental data

The study was conducted at the Santa Lucia Foundation IRCCS of Rome, in accordance with the ethical standards of the local institutional review board and with the 1964 Helsinki declaration and its later amendments. The study protocol was approved by the local ethics committee (Approval 2022_010), and all participants provided written informed consent prior to enrollment. For this study, 37 healthy elderly subjects were initially recruited, but only 33 participants (18 females, age 65.5 ± 7.5 years, mean ± SD) completed the protocol (see below). Exclusion criteria included the presence of neurological or psychiatric disorders, significant cardiovascular or cerebrovascular disease, uncontrolled hypertension, diabetes with end-organ damage, and contraindications to MRI. All subjects underwent two MRI sessions on a Siemens MAGNETOM 3T Prisma scanner. An in-house built MR compatible gas delivery system was used for the respiratory paradigm, which consisted in alternating between medical air (<0.04% CO_2_, 21% O_2_, 78% N_2_, ∼1% noble gases) and a hypercapnic gas mixture (5% CO_2_, 21% O_2_, 74% N_2_) every 2 min across two 10-min sessions. The first session included a PASL-fMRI sequence (for CBF) with voxel size of 2.2×2.2×3 mm^3^ nominal resolution, TR 5.5 s (for pair control-label), TE 13.26 ms, 110 volumes. The second session included a VASOBOLD-fMRI (for combined CBV and BOLD) with voxel size of 2.2×2.2×3 mm^3^ nominal resolution, TR 2.7 s, TE 14.3 ms, inversion delay 550 ms, 224 paired volumes. All timeseries were denoised using NORDIC (Vizioli et al., 2021), realigned for motion with ASLtbx (Wang et al., 2008), split from the BOLD contrast, and corrected for BOLD contamination using LAYNII (Huber et al., 2021). Structural images (T1-weighted MEMPRAGE, 1.0×1.0×1.0 mm^3^) were segmented into gray matter, white matter, and cerebrospinal fluid using the CAT12 (Gaser et al., 2024) segmentation tool. The BOLD and VASO functional timeseries were processed using seeVR (Bhogal, 2021). Maps of CBF, CBV, and BOLD were spatially normalized to the MNI 2 mm template using ANTs (Tustison et al., 2021). Partial pressures of end-tidal O_2_ (p_ET_O_2_) and CO_2_ (p_ET_CO_2_) were extracted from raw capnography data as the lower and upper envelopes, respectively, and the resulting temporal traces were assessed for quality. Subjects with poor capnography data (3 subjects) and/or incomplete data acquisition (1 subject) were excluded from further analysis.

### Data-driven model optimization of vascular parameters

To characterize the temporal dynamics of the cerebrovascular response to hypercapnia, we extracted three complementary parameters, namely delay, time constant, and gain, mapping the partial pressure of end-tidal gases (p_ET_O_2_ and p_ET_CO_2_) to the arterial values in the brain (PaO_2_ and PaCO_2_). The delay (*δ*) was defined as the interval between the onset of the hypercapnic challenge (corrected for instrumental gas delivery delays and analyzer lag) and the resulting rise in regional arterial capillary gases. This operational definition aligns with commonly used metrics of vascular response latency in dynamic cerebrovascular reactivity (CVR) studies, where delay represents the temporal lag between the respiratory challenge and the onset of the vascular response (Poulin et al., 1996; Tancredi & Hoge, 2013). The time constant (*τ*) was obtained from an exponential kernel used to convolve the end-tidal gases time course with the vascular response. In this context, the time constant quantifies the rate at which the regional arterial capillary partial pressure approaches its new steady-state following a change in inhaled gases, providing a measure of the system’s dynamic responsiveness. This approach is consistent with first-order linear system models commonly employed in literature, where *τ* reflects the intrinsic kinetics of the cerebrovascular response (Tancredi & Hoge, 2013). The gain (*κ*) was obtained from scaling the incremental portion of the vascular response. In our framework, delay, time constant, and gain are not applied directly to the relationship between end-tidal gases and CBF. Instead, they are used to model the transfer from end-tidal to arterial gases (ℎ_*pET*→*Pa*_), which in turn influence cerebrovascular tone. These parameters thus describe the fidelity and temporal accuracy with which respiratory gas variations are transmitted into the cerebral circulation, capturing the initiation, rate, and magnitude of vascular adaptation. This approach provides complementary information to conventional cerebrovascular reactivity (CVR) metrics.

We extracted vascular parameters in a region-of-interest (ROI) based manner, using a commonly employed 400 parcel human parcellation atlas (Schaefer et al., 2018). For each ROI, we used the model to simulate CBF response to hypercapnia using the experimental basal flow (CBF_0_, averaged from the first 21 PASL-fMRI volumes, corresponding to 2 minutes of resting control conditions) and hypercapnic data (p_ET_O_2_ and p_ET_CO_2_). We adjusted the delay, time constant, and gain of the ℎ_*pET*→*Pa*_(*δ*, *τ*, *κ*) transfer-function iteratively using a sign-based stochastic gradient descent (SGD) algorithm (32 iterations; initial step sizes: 0.4 s and 0.8 s for *δ* and *τ*, respectively; decay rate: 0.1). The objective function for minimization was defined as the residual sum of squares (RSS) between simulated and experimental CBF. The latter was used to update *κ* at each step using linear regression. Optimization parameters were preliminarily tuned using a 16×16 (*δ*×*τ*) grid search (with *κ* unconstrained), which provided the boundaries of *δ* (0-10 s), *τ* (0-24 s), and *κ* (0-2) as well as appropriate global (i.e., equal for all ROIs) uniform priors as domains for the GSD-based initialization (1-7 s for *δ*, 4-16 s for *τ*, and 0.5-1.5 for *κ*). Since the model accepts CBF_0_ values in the range 15-180 mL/100g/min, voxels with mean CBF_0_ falling outside this range (<10%) were excluded from the simulations. All ROIs in all subjects contained >90% of valid voxels.

### Statistical analysis

All statistical analyses were performed in MATLAB (R2025a) using the Statistics and Machine Learning Toolbox. Parameter uncertainty for the fitted transfer-function parameters 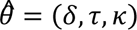 was quantified using the asymptotic covariance matrix derived from the Jacobian *J* (Gauss-Newton approximation) of the model output at the optimum (i.e., with respect to parameters at the fitted solution). Numerically, we computed finite-difference sensitivities of the simulated CBF to small perturbations of each parameter, estimated the residual variance 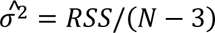, where *N* is the number of MRI volumes, and formed 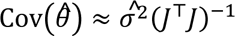. The standard error (SE) for each parameter estimate was calculated as the square root of the diagonal elements of the covariance matrix, and 95% confidence intervals (CIs) were derived as 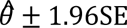. For robustness, we used model fit-quality metrics to down-weight or exclude poorly fitted ROIs in subsequent analyses. To assess distributional assumptions, the normality of parameter distributions across subjects was tested ROI-wise using both the Lilliefors and Shapiro-Wilk tests. All extracted vascular parameters conformed to normality in >60% of ROIs, and therefore they were analyzed with parametric methods.

Group-level inference proceeded in two steps. First, linear mixed-effects models were fitted separately for each parameter to test whether values differed across ROIs, with ROI as a fixed effect and subject as a random effect. Observation-level weights equal to the inverse variance (1/SE^2^) were used to account for heteroscedasticity of the estimates. Omnibus ROI effects were assessed by Satterthwaite-corrected ANOVA of the mixed model. When required, post-hoc comparisons were adjusted for multiple testing using false discovery rate (FDR) correction. Second, associations with age (range 51-80 years) were examined. Weighted linear mixed-effects models were fitted with parameter value as the dependent variable, age (and sex as covariate) as fixed effects, and random intercepts for subject and ROI. This tested for global age effects across all ROIs. In addition, ROI-wise weighted linear regressions were performed to localize age-related effects. Resulting *p*-values were corrected for multiple comparisons using FDR. All tests were two-tailed with nominal *α*=0.05 after correction.

### Code and data availability

The code utilized in this study is accessible on GitHub ^2^. Data is available from the corresponding author upon reasonable request and completion of required forms ^3^.

## Results

### Model validation

Model simulations reproduced several established neurovascular and neurometabolic relationships reported in the literature (see also the Materials and methods section). The model captured the well-known coupling between neurotransmitter cycling and neuronal oxidative glucose consumption as well as the association between incremental changes in CMRO_2_ and CBF observed during activation (**Figure 1**). In both cases, the simulated steady-state showed a slightly non-linear trend compared with the typically linear fits commonly applied to human data (but see Hyder et al., 2002; Rothman et al., 2022), however the overall agreement was robust, and simulated data remained within the 95% prediction interval. Beyond steady-state relations, the model reproduced the stereotypical temporal dynamics of neuroimaging and metabolic signals during different levels of simulated brain activation (**Figure 2**) and during simulated graded hypercapnia (**Figure 3**). Finally, the model accurately predicted cerebrovascular reactivity (CVR) to hypercapnia, showing that BOLD-, CBF-, and CBV-derived indices converged on the same CVR values regardless of the magnitude of the hypercapnic challenge (**Figure 4**). Taken together, these validation results confirm that the model captures key neurovascular and neurometabolic relationships across both steady-state and dynamic conditions, providing a reliable framework for the analyses that follow.

**Figure 1.**
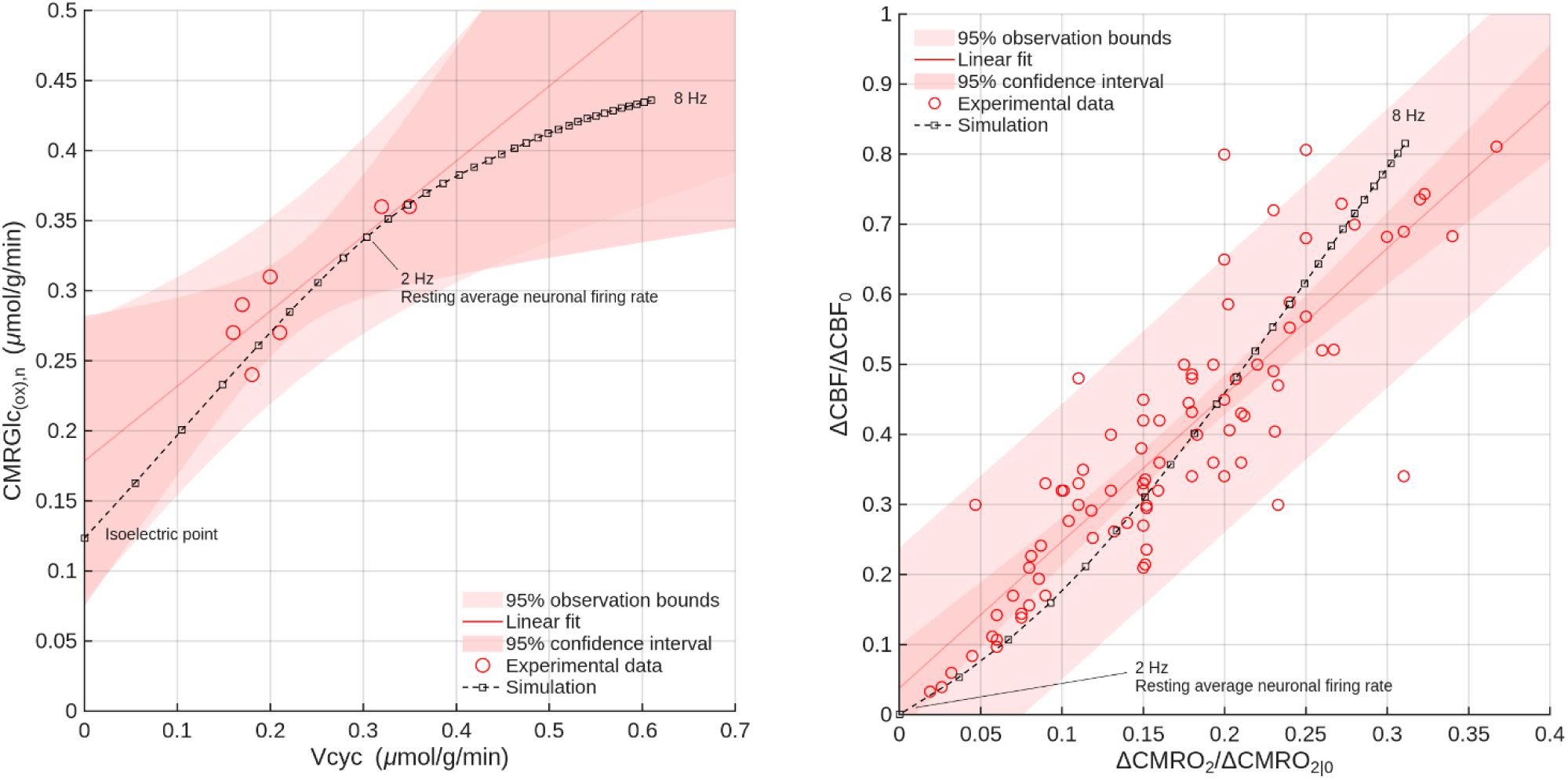
Model validation against steady-state relationships. *Left*: Simulated relationship between neurotransmitter cycling rate (V_cyc_) and neuronal oxidative glucose consumption (CMRGlc_(ox),n_, i.e., neuronal CMRO_2_). Human experimental data are sparse outside the resting state, but simulated values (solid line, 95% CI shaded) fell within the confidence bounds of reported measurements. *Right*: Simulated relationship between incremental changes in CMRO_2_ and CBF across a wide range of neuronal activity levels. Model predictions exhibited a slightly non-linear trend, while experimental data are typically fitted linearly. Residuals confirmed that deviations were small and systematic (not shown). All experimental data were derived from our recently published meta-analysis (Rothman et al., 2022).

**Figure 2.**
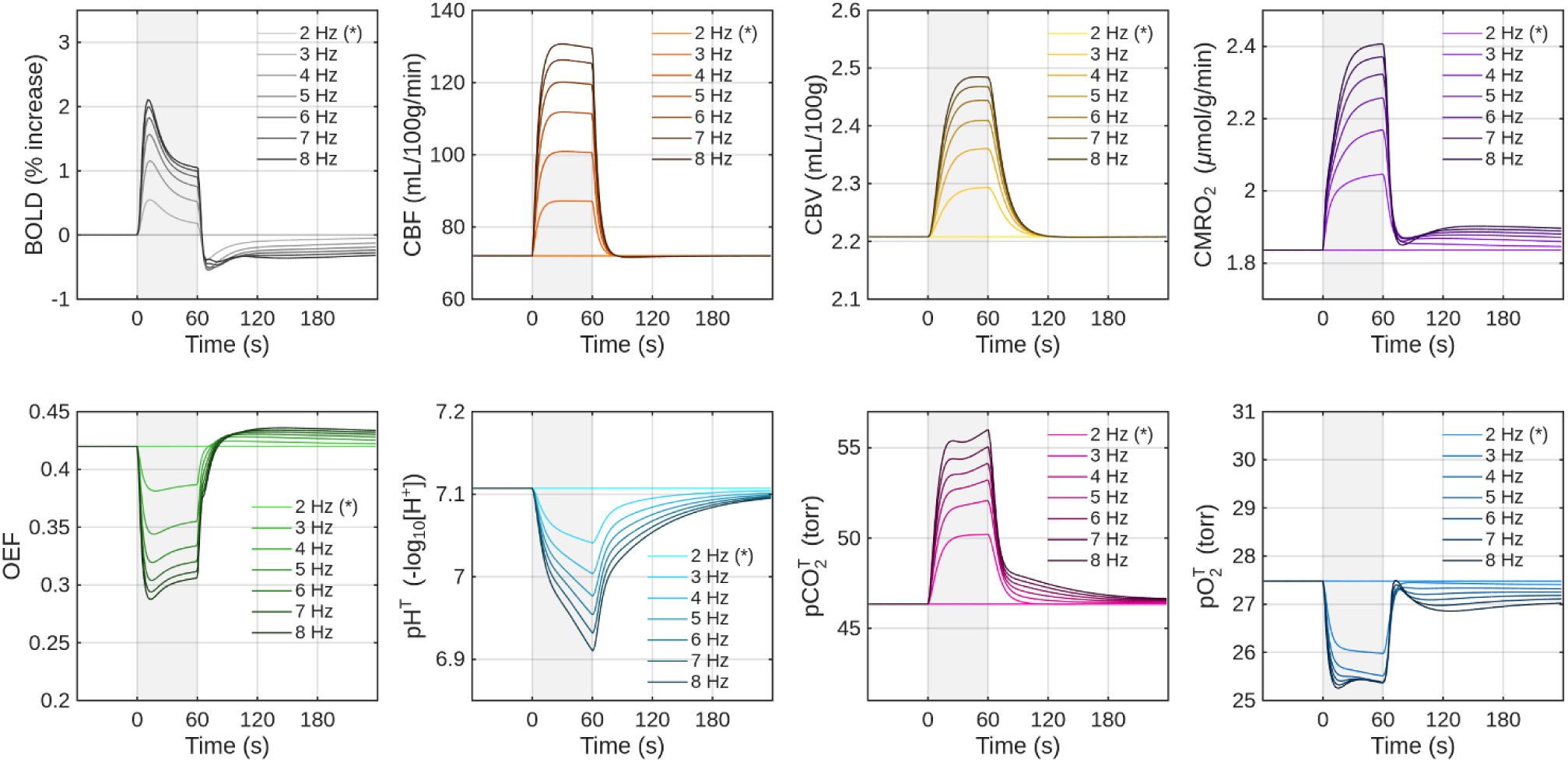
Simulated responses to graded neuronal activation. Predicted time courses of BOLD, CBF, CBV, CMRO_2_, OEF, and tissue gas variables (pH, pCO_2_, pO_2_) during a simulated graded neuronal activation (2-8 Hz firing rates, corresponding to V_cyc_ values in the approx. range 0.3-0.6 µmol/g/min). Simulated dynamics were stereotypical and consistent with reported experimental findings. During simulated graded neuronal activation, predicted responses of BOLD, CBF, CBV, CMRO_2_, OEF, and tissue gases (pH, pCO_2_, pO_2_) were in good agreement with reported experimental findings or calculations derived from meta-analyses and mathematical modeling (DiNuzzo et al., 2024).

**Figure 3.**
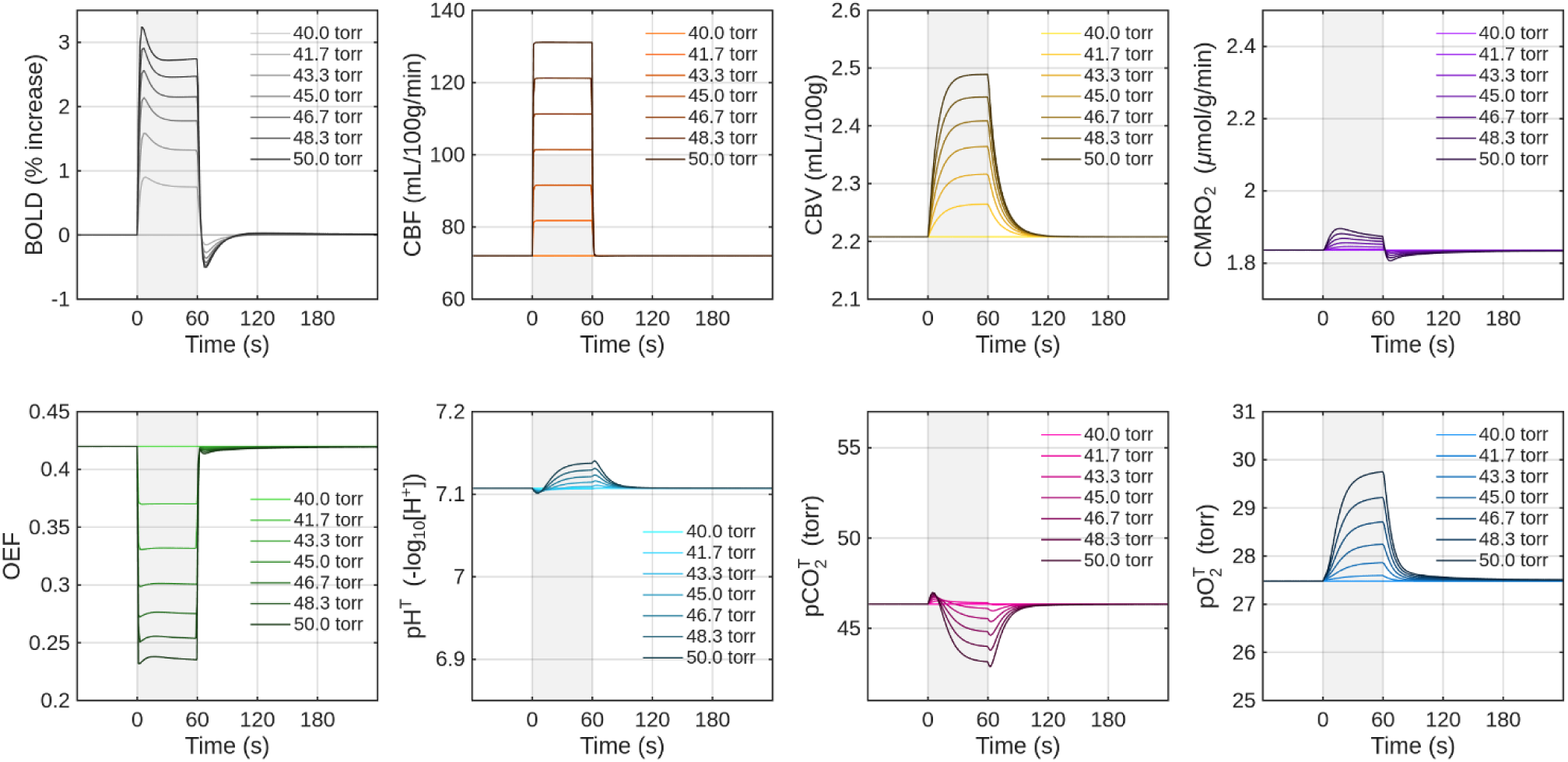
Simulated responses to graded hypercapnia. Predicted time courses of BOLD, CBF, CBV, CMRO_2_, OEF, and tissue gas variables (pH, pCO_2_, pO_2_) during a simulated increase in arterial pCO_2_ from 40 to 50 mmHg. Responses reproduced the characteristic hypercapnia-induced vascular and metabolic changes described in the literature. For graded hypercapnic challenge (from a baseline of 40 mmHg up to 50 mmHg), the model yielded the characteristic increases in BOLD, CBF, and CBV, along with the expected shifts in metabolic variables derived from meta-analyses and mathematical modeling (DiNuzzo et al., 2024).

**Figure 4.**
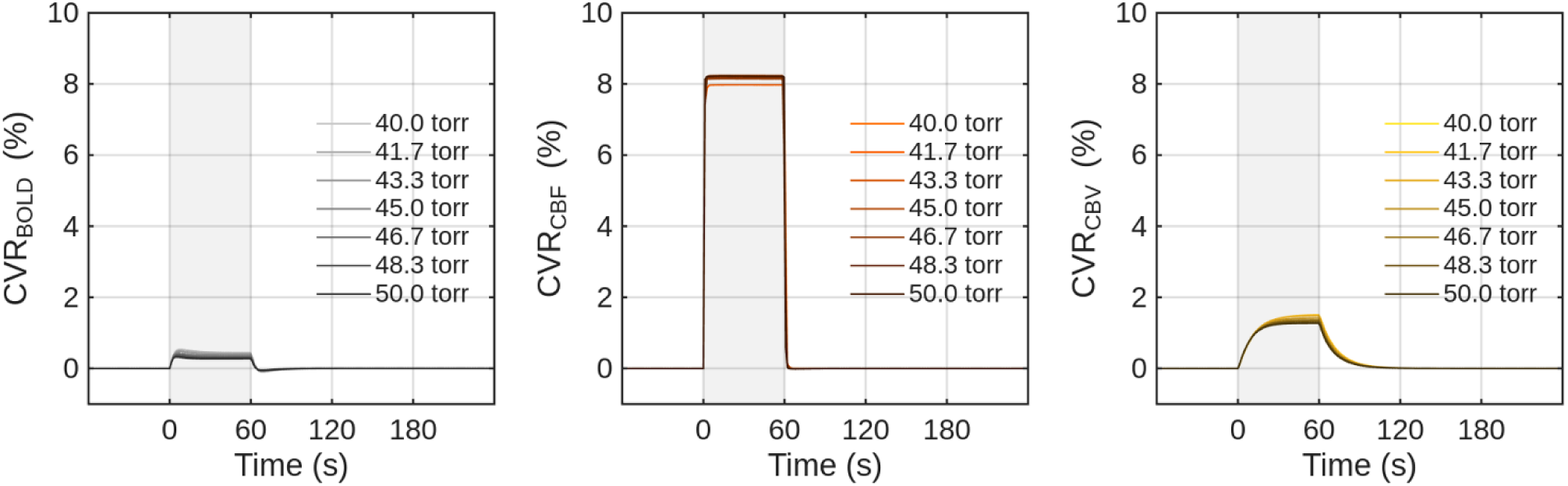
Prediction of cerebrovascular reactivity (CVR). Simulated CVR derived from BOLD, CBF, and CBV signals during graded hypercapnia. Simulations for all three modalities converged on the same CVR values regardless of the level of hypercapnic challenge, in line with experimental observations. This independence from stimulus strength mirrors experimental observations, strengthening the model’s validity.

### Parameter estimation and distributions

We next applied the model to experimental CBF data in our cohort, estimating subject-specific cerebrovascular parameters with quantified uncertainty. Model fitting to the ASL-derived CBF time courses converged for all 33 subjects and 400 cortical ROIs, and provided subject-specific estimates of vascular parameters, with SE and CI quantified by the asymptotic covariance of the Jacobian (see Materials and methods section). These estimates were used as inverse-variance weights in subsequent analyses. Across subjects, we obtained a mean delay (*δ*) of 5.23 ± 1.44 s (mean ± SD, group 95% CI: 4.72-5.74 s), a mean time constant (*τ*) of 9.78 ± 3.91 s (mean ± SD, group 95% CI: 8.40-11.17 s), and a mean gain (*κ*) of 0.96 ± 0.16 (mean ± SD, group 95% CI: 0.90-1.01) (see **Supplementary Table 5** for the results in all 400 ROIs). These estimates indicate that cerebrovascular responses in healthy aging are in the order of seconds for onset and adaptation, with preserved overall amplitude of response. The distributions of key parameters were consistent across subjects, though individual variability was evident. In representative subjects, posterior parameter distributions showed narrow confidence intervals, confirming stable optimization, while in few subjects the distributions were broad, apparently bimodal, and/or skewed, which might reflect changes limited to specific brain regions (**Figure 5**). Across the cohort, the vascular parameters exhibited nonsignificant correlations. In particular, delay (*δ*) and time constant (*τ*) were weakly positively associated (*r* = 0.20, *p* = 0.68), whereas gain (*κ*) appeared largely independent of both *δ* (*r* = −0.05, *p* = 0.32) and *τ* (*r* = 0.02, *p* = 0.76).

**Figure 5.**
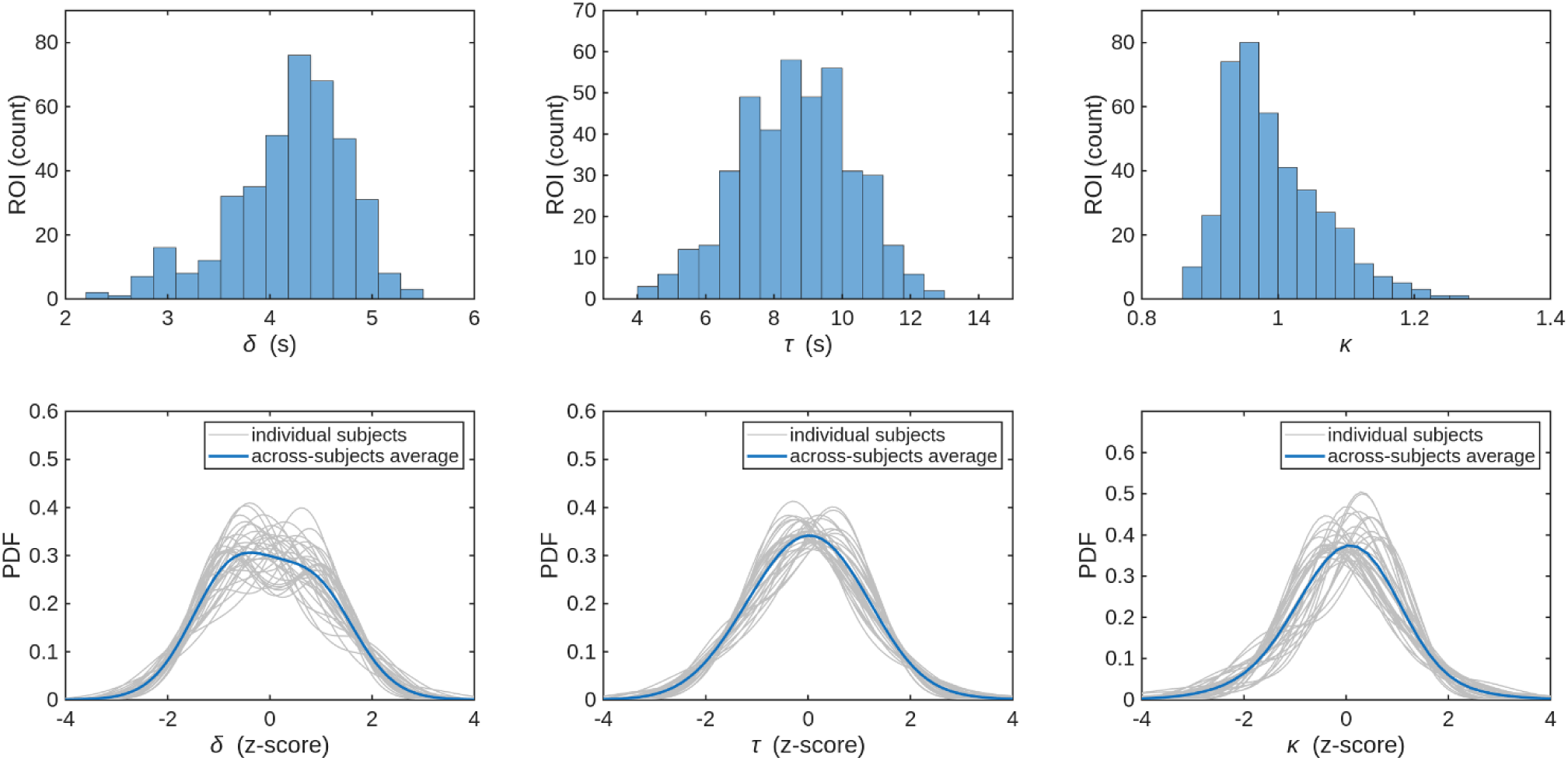
Distributions of estimated vascular parameters. Posterior distributions of the optimized vascular parameters obtained from ASL-derived CBF fitting. *Top row*: histogram from a representative subject showing the uncertainty intervals around the posterior maxima. *Bottom row*: group-level probability density function (PDF) derived from kernel density estimation across all subjects, indicating consistent parameter convergence. Significant departures from normality are apparent in some subjects, especially for *δ*, where apparent bimodality is observed in some subjects.

### Cross-modality validation against independent measurements

To assess the predictive value of the estimated parameters, we simulated CBF, CBV and BOLD responses using the optimized parameter sets. The simulated CBF responses closely matched the ASL data used for fitting, while the model also reproduced CBV and BOLD dynamics measured independently with VASO during a separate imaging session, when provided with concurrent CBF₀ and end-tidal gas traces (**Figure 6**). This agreement supports the internal consistency of the model and its ability to generalize across modalities.

**Figure 6.**
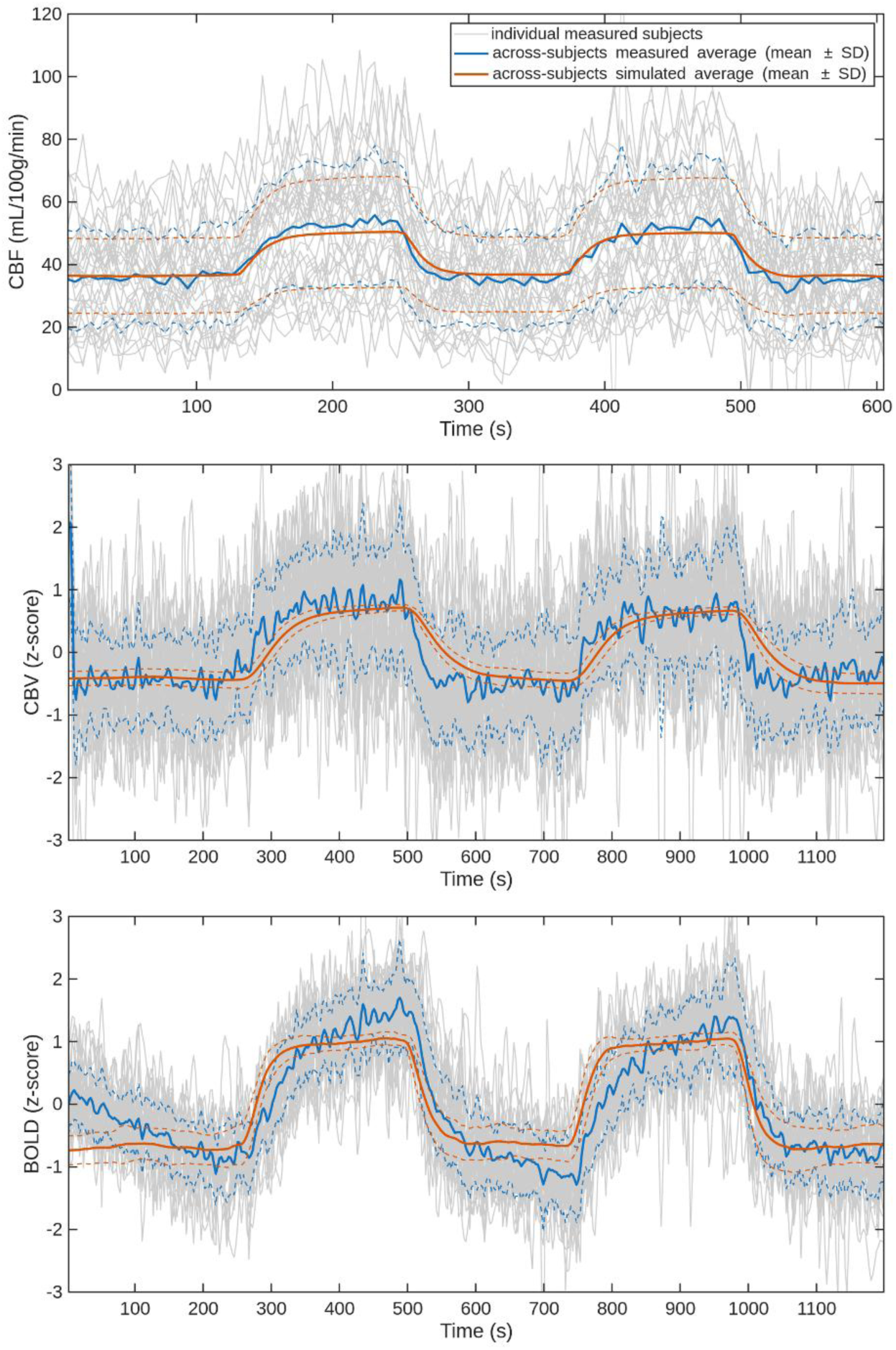
Validation of model predictions across modalities. Simulated time courses of CBF (quantitative, fitted to ASL), CBV, and BOLD (nonquantitative, predicted and compared with VASO data acquired in a separate session). When parametrized with subject-specific values and provided with concurrent baseline CBF₀ and end-tidal gas traces, the model reproduced both training (ASL) and independent (VASO) measurements with good fidelity.

### Spatial distribution of vascular parameters

The optimized parameters were then mapped voxel-wise. Group-level average maps highlighted systematic differences across brain areas (**Figure 7**), reflecting the substantial regional heterogeneity in individual subjects. We evaluated whether vascular parameters varied across brain regions using linear mixed-effects models with ROI as a fixed effect and subject as a random effect (**Table 1** and **Supplementary Table 6**). The omnibus tests revealed highly significant ROI effects for all three parameters. Specifically, for *δ* the ROI effect was significant (F_(1,13171)_ = 73.08, *p* < 1.4×10^-17^), indicating systematic differences in onset latency across regions. Similarly, *τ* showed a significant ROI effect (F_(1,13167)_ = 34.22, *p* < 5.0×10^-9^), reflecting regional variability in the rate of vascular adjustment. Finally, *κ* also exhibited a significant, though smaller, ROI effect (F_(1,13168)_ = 14.79, *p* = 1.2×10^-4^), again suggesting that the amplitude of the CO_2_-driven response also varies between regions. Overall, these findings indicate that cerebrovascular dynamics are not uniform across the brain. A complementary analysis slicing the brain along the left-right, anterior-posterior, and superior-inferior axes confirmed subtle asymmetries at the subject-level, yet an overall stability at group-level (**Figure 8**).

**Figure 7.**
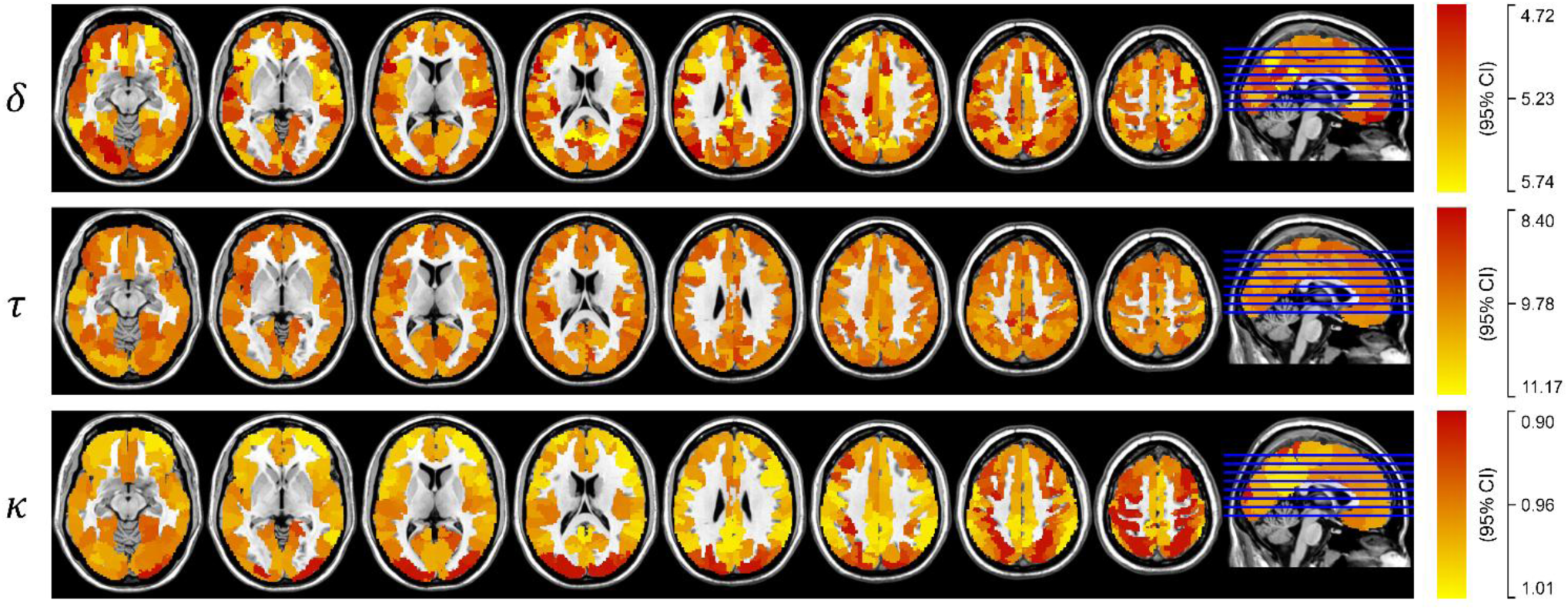
Spatial distribution of vascular parameters. Group averaged voxel-wise maps of optimized vascular parameters (*top*: *δ*, *middle*: *τ*, *bottom*: *κ*). Regional heterogeneity across specific cortical areas is apparent, reflecting the substantial inter-subject heterogeneity.

**Figure 8.**
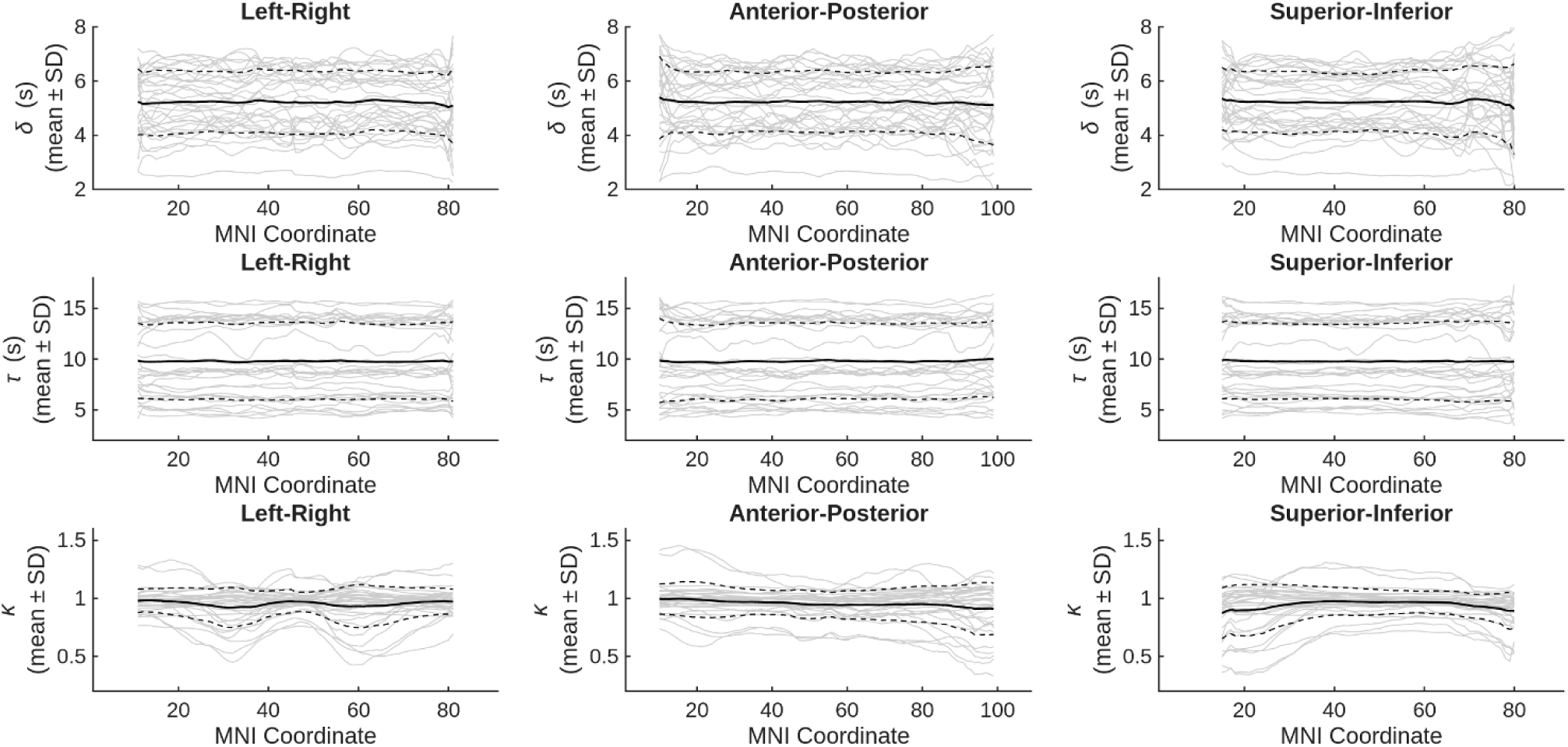
Parameter distributions in vascular territories and spatial gradients. Sliced analysis of vascular parameters (*top*: *δ*, *middle*: *τ*, *bottom*: *κ*) along the left-right (*left*), anterior-posterior (*center*), and superior-inferior axes (*right*), showing the absence of prominent asymmetries at the macroscopic-scale, in spite of the high degree of variability at the single-subject level.

**Table 1.** Spatial variability of optimized cerebrovascular parameters across ROIs in healthy aging adults. Summary statistics across the 10 most significant (FDR-corrected *p*-value) ROIs.

| ROI | Mean | SD | Effect Size | *p*-value | Direction |
| --- | --- | --- | --- | --- | --- |
| *Delay ($\delta$)* |  |  |  |  |  |
| LH_SomMotB_Aud_2 | 5.010086 | 1.669609 | 3.000755 | 0 | Below Mean |
| LH_TempPar_2 | 4.726288 | 1.733381 | 2.726629 | 0 | Below Mean |
| RH_SalVentAttnB_PFCI_3 | 4.65009 | 1.581663 | 2.94 | 0 | Below Mean |
| RH_ContA_PFCI_1 | 5.431155 | 1.936146 | 2.805136 | 4.44E-14 | Above Mean |
| RH_ContC_pCun_4 | 5.365811 | 1.852798 | 2.896058 | 4.62E-13 | Above Mean |
| LH_DefaultA_PFCd_3 | 4.984002 | 1.732126 | 2.877391 | 6.96E-13 | Below Mean |
| RH_ContB_IPL_4 | 5.317129 | 1.693555 | 3.139625 | 1.81E-10 | Above Mean |
| LH_SomMotB_S2_5 | 5.508306 | 1.875191 | 2.937464 | 5.34E-09 | Above Mean |
| RH_SalVentAttnA_FrMed_3 | 5.058186 | 1.816742 | 2.784207 | 4.21E-08 | Below Mean |
| RH_SomMotA_4 | 5.416448 | 2.088457 | 2.593516 | 4.43E-08 | Above Mean |
| *Time constant ($\tau$)* |  |  |  |  |  |
| LH_VisCent_ExStr_1 | 0.981808 | 0.097811 | 10.03786 | 0 | Above Mean |
| LH_VisPeri_ExStrInf_1 | 0.95373 | 0.080419 | 11.8595 | 0 | Below Mean |
| LH_VisPeri_ExStrInf_2 | 0.963752 | 0.071401 | 13.4978 | 0 | Above Mean |
| LH_VisPeri_ExStrInf_3 | 0.974329 | 0.078285 | 12.44591 | 0 | Above Mean |
| LH_VisPeri_ExStrInf_4 | 0.946127 | 0.08779 | 10.7772 | 0 | Below Mean |
| LH_VisPeri_ExStrInf_5 | 0.949804 | 0.068761 | 13.81313 | 0 | Below Mean |
| LH_VisPeri_StriCal_1 | 0.977566 | 0.134275 | 7.280321 | 0 | Above Mean |
| LH_VisPeri_StriCal_2 | 0.977601 | 0.068327 | 14.30774 | 0 | Above Mean |
| LH_VisPeri_ExStrSup_1 | 0.955945 | 0.057278 | 16.68969 | 0 | Below Mean |
| LH_VisPeri_ExStrSup_2 | 0.977891 | 0.141003 | 6.935254 | 0 | Above Mean |
| *Gain ($\kappa$)* |  |  |  |  |  |
| LH_VisPeri_ExStrSup_1 | 10.16754 | 4.375973 | 2.323493 | 0 | Above Mean |
| LH_DefaultB_PFCv_2 | 9.560725 | 4.014846 | 2.381343 | 0 | Below Mean |
| LH_TempPar_2 | 9.989894 | 4.005952 | 2.493763 | 0 | Above Mean |
| RH_ContB_IPL_4 | 10.21132 | 4.107677 | 2.485912 | 4.44E-13 | Above Mean |
| RH_ContC_pCun_3 | 10.09266 | 3.887932 | 2.595895 | 3.77E-10 | Above Mean |
| RH_SalVentAttnA_PrC_1 | 9.597167 | 4.364506 | 2.198913 | 1.44E-09 | Below Mean |
| LH_DorsAttnB_PostC_2 | 9.743768 | 4.163909 | 2.340053 | 7.06E-09 | Below Mean |
| RH_VisCent_ExStr_1 | 9.176746 | 3.952695 | 2.321643 | 1.03E-08 | Below Mean |
| RH_DorsAttnA_TempOcc_2 | 9.771381 | 4.252401 | 2.297851 | 1.03E-08 | Below Mean |
| RH_DefaultB_PFCd_3 | 10.13677 | 4.362254 | 2.323746 | 5.90E-08 | Above Mean |

### Simulated CMRO_2_ and tissue pH, pO_2_, and pCO_2_

From the optimized vascular parameters, model simulations of CMRO_2_ confirmed that the variation of oxidative metabolism during hypercapnia was minimal across subjects, in line with the prevailing interpretation of metabolic stability under this condition (**Figure 9**). At the same time, simulated tissue pH revealed small but consistent acidifications. These pH shifts are quantitatively within the range reported by recent studies suggesting that even small acid-base disturbances may contribute to functional brain impairments.

**Figure 9.**
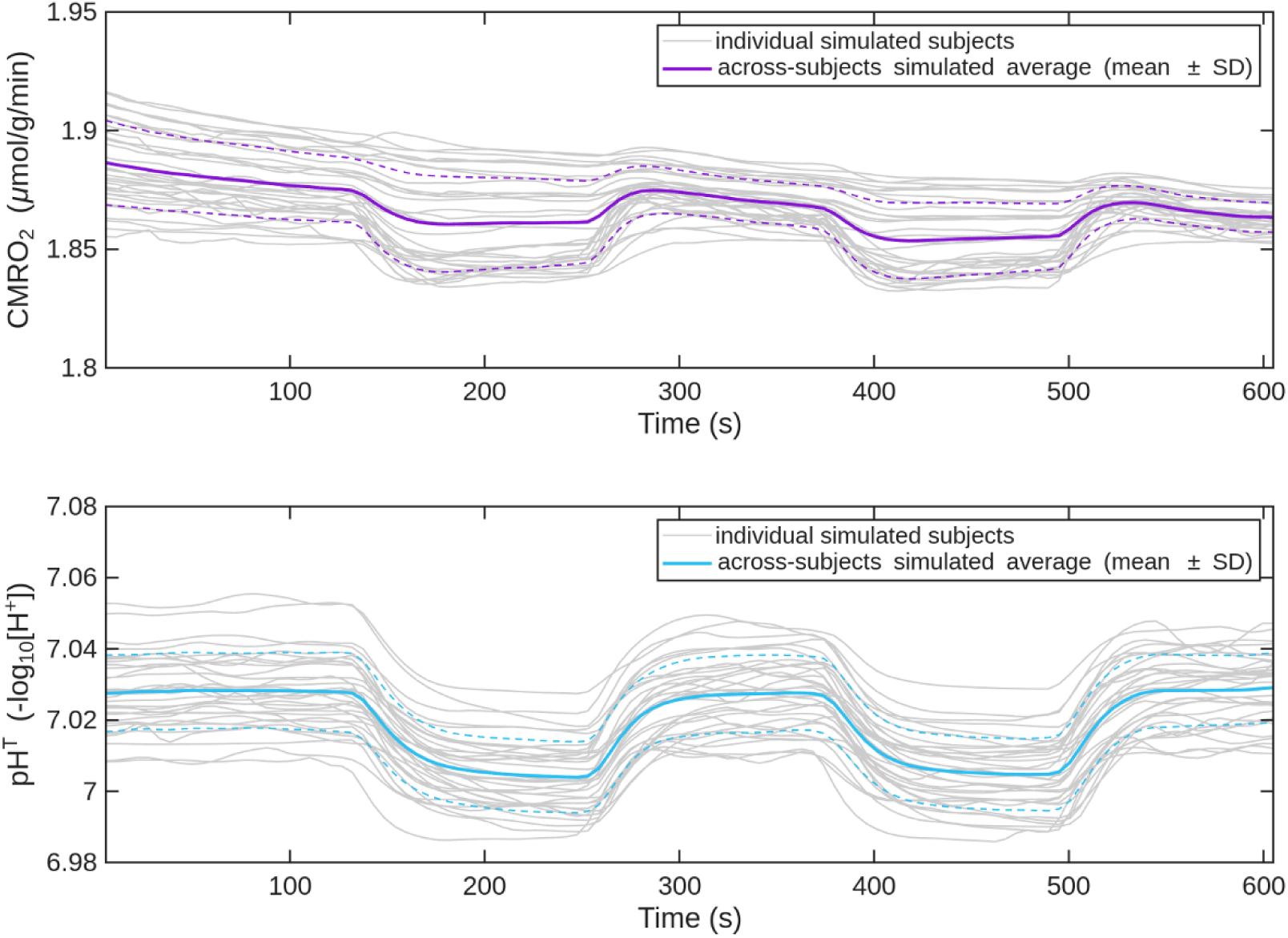
Simulated CMRO₂ and tissue pH responses. *Top*: Predicted CMRO_2_ changes (around 1% on average) across subjects during hypercapnia, confirming minimal variability and stability of oxidative metabolism. *Bottom*: Predicted tissue (i.e., volume-weighted average in cellular and extracellular compartments) pH changes, showing subtle (∼0.03 pH units) yet possibly physiologically relevant acidification.

### Age dependence of vascular parameters

Finally, we examined the association between vascular parameters and age using linear mixed-effects models, with subject as a random effect and Age and Sex as fixed effects (**Table 2** and **Supplementary Table 7**). Across the whole brain, we found no significant age-related changes in *δ* (F_(1,32.4)_ = 0.65, *p* = 0.43) or *τ* (F_(1,32.0)_ = 0.09, *p* = 0.76). In contrast, *κ* showed a modest but significant decrease with age (F_(1,32.8)_ = 5.08, *p* = 0.031), suggesting that the overall amplitude of the CO_2_ response tends to diminish in older subjects. Sex did not significantly influence *δ* or *τ*, while a small effect was observed for *κ* (*p* = 0.046). At the ROI level, FDR-corrected analyses revealed that age was significantly associated with *δ* in 68 out of 400 ROIs and with *τ* in 67 ROIs, indicating region-specific subtle alterations of vascular dynamics with aging. Instead, *κ* exhibited a stronger and more widespread age effect, with 320 out of 400 ROIs showing significant age-related reductions, consistent with the global trend observed in the mixed-effects model. It should be noted however that across the cohort, age was associated with heterogeneous changes in vascular parameters. For *δ*, several regions exhibited increases with age (slower response dynamics), whereas other regions showed decreases. For *τ* and κ, most significant effects were negative (reduced time constant and reduced gain with age), but a subset of ROIs still showed the opposite trend.

**Table 2.**
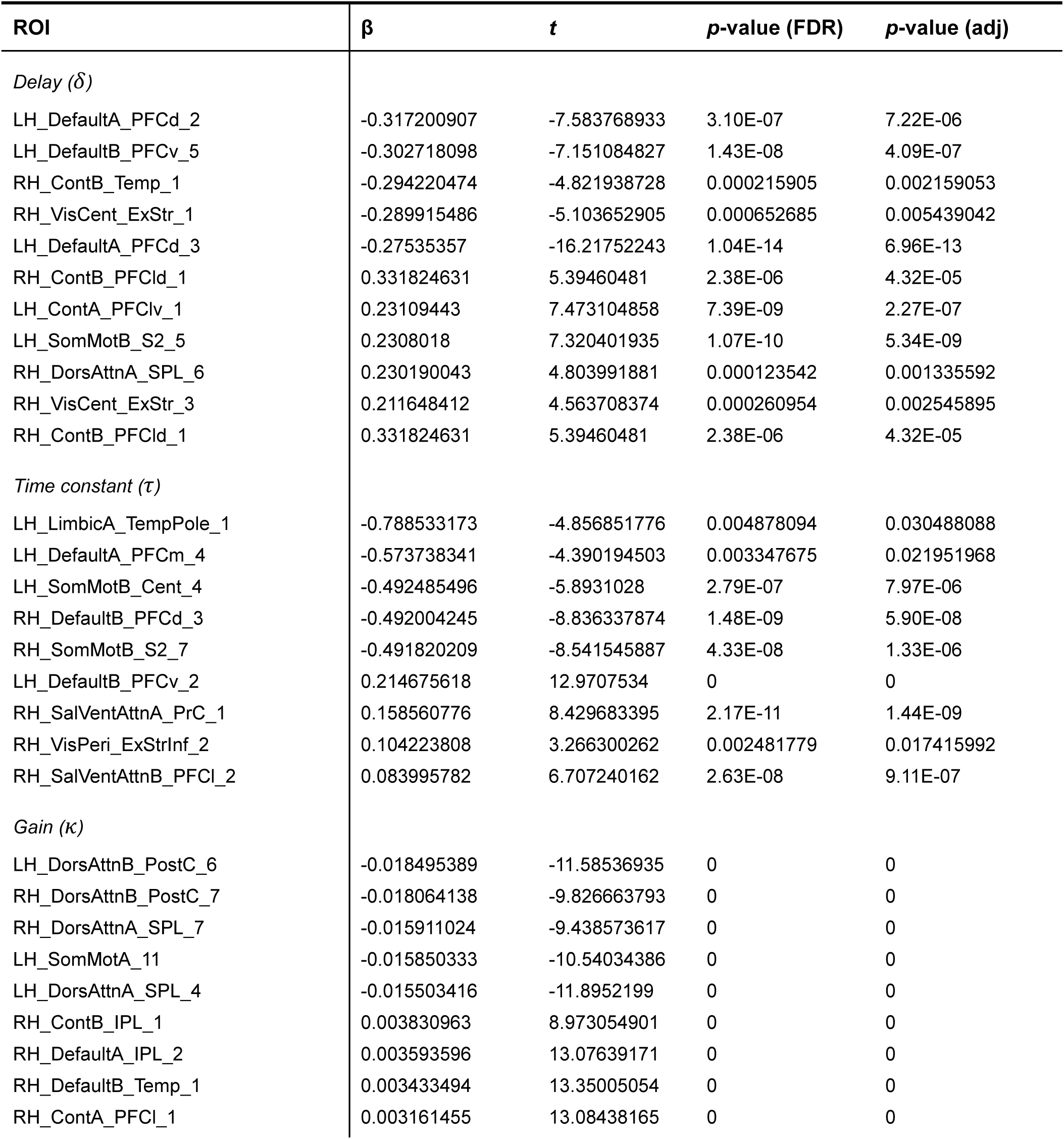
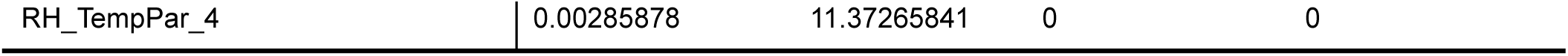
Age-related associations of optimized cerebrovascular parameters across ROIs in healthy aging adults. Summary statistics across the 10 most significant (FDR-corrected and Benjamini-Hochberg adjusted *p*-values) ROIs (top 5 for β < 0 and top 5 for β > 0).

| ROI | $\beta$ | $t$ | $p$-value (FDR) | $p$-value (adj) |
| --- | --- | --- | --- | --- |
| *Delay ($\delta$)* |  |  |  |  |
| LH_DefaultA_PFCd_2 | -0.317200907 | -7.583768933 | 3.10E-07 | 7.22E-06 |
| LH_DefaultB_PFCv_5 | -0.302718098 | -7.151084827 | 1.43E-08 | 4.09E-07 |
| RH_ContB_Temp_1 | -0.294220474 | -4.821938728 | 0.000215905 | 0.002159053 |
| RH_VisCent_ExStr_1 | -0.289915486 | -5.103652905 | 0.000652685 | 0.005439042 |
| LH_DefaultA_PFCd_3 | -0.27535357 | -16.21752243 | 1.04E-14 | 6.96E-13 |
| RH_ContB_PFCId_1 | 0.331824631 | 5.39460481 | 2.38E-06 | 4.32E-05 |
| LH_ContA_PFCIv_1 | 0.23109443 | 7.473104858 | 7.39E-09 | 2.27E-07 |
| LH_SomMotB_S2_5 | 0.2308018 | 7.320401935 | 1.07E-10 | 5.34E-09 |
| RH_DorsAttnA_SPL_6 | 0.230190043 | 4.803991881 | 0.000123542 | 0.001335592 |
| RH_VisCent_ExStr_3 | 0.211648412 | 4.563708374 | 0.000260954 | 0.002545895 |
| RH_ContB_PFCId_1 | 0.331824631 | 5.39460481 | 2.38E-06 | 4.32E-05 |
| *Time constant ($\tau$)* |  |  |  |  |
| LH_LimbicA_TempPole_1 | -0.788533173 | -4.856851776 | 0.004878094 | 0.030488088 |
| LH_DefaultA_PFCm_4 | -0.573738341 | -4.390194503 | 0.003347675 | 0.021951968 |
| LH_SomMotB_Cent_4 | -0.492485496 | -5.8931028 | 2.79E-07 | 7.97E-06 |
| RH_DefaultB_PFCd_3 | -0.492004245 | -8.836337874 | 1.48E-09 | 5.90E-08 |
| RH_SomMotB_S2_7 | -0.491820209 | -8.541545887 | 4.33E-08 | 1.33E-06 |
| LH_DefaultB_PFCv_2 | 0.214675618 | 12.9707534 | 0 | 0 |
| RH_SalVentAttnA_PrC_1 | 0.158560776 | 8.429683395 | 2.17E-11 | 1.44E-09 |
| RH_VisPeri_ExStrInf_2 | 0.104223808 | 3.266300262 | 0.002481779 | 0.017415992 |
| RH_SalVentAttnB_PFCI_2 | 0.083995782 | 6.707240162 | 2.63E-08 | 9.11E-07 |
| *Gain ($\kappa$)* |  |  |  |  |
| LH_DorsAttnB_PostC_6 | -0.018495389 | -11.58536935 | 0 | 0 |
| RH_DorsAttnB_PostC_7 | -0.018064138 | -9.826663793 | 0 | 0 |
| RH_DorsAttnA_SPL_7 | -0.015911024 | -9.438573617 | 0 | 0 |
| LH_SomMotA_11 | -0.015850333 | -10.54034386 | 0 | 0 |
| LH_DorsAttnA_SPL_4 | -0.015503416 | -11.8952199 | 0 | 0 |
| RH_ContB_IPL_1 | 0.003830963 | 8.973054901 | 0 | 0 |
| RH_DefaultA_IPL_2 | 0.003593596 | 13.07639171 | 0 | 0 |
| RH_DefaultB_Temp_1 | 0.003433494 | 13.35005054 | 0 | 0 |
| RH_ContA_PFCI_1 | 0.003161455 | 13.08438165 | 0 | 0 |
| RH_TempPar_4 | 0.00285878 | 11.37265841 | 0 | 0 |

## Discussion

Our previous modeling efforts have provided a mechanistic foundation for interpreting cerebrovascular responses in both health and disease. Specifically, we highlighted that neurovascular coupling (NVC) is not solely tuned to deliver oxygen, but also to remove protons and CO_2_, thereby maintaining homeostasis of pH, pCO_2_, and pO_2_ (DiNuzzo et al., 2024). This work suggested that any alterations in CBF dynamics, whether in amplitude or temporal characteristics, could disrupt tissue homeostasis and impact brain function. Building on this concept, we recently used our modeling framework in the context of aging, showing that age-associated reductions in regional CBF, even in the presence of largely preserved oxygen consumption, lead to local increases in pCO_2_ and acid shifts in tissue pH (Mangia et al., 2025). These metabolic perturbations have plausible functional consequences and provide a mechanistic link between declining vascular health and early cognitive alterations.

The current study further extends these insights by quantifying dynamic features of the CBF response in healthy aging. While our previous work focused on steady-state or integrated measures of vascular and metabolic balance, the present analysis addresses the temporal responsiveness of the cerebrovascular system to a physiological challenge (hypercapnia). By demonstrating progressive alteration in delay, time constants, and gain of the vascular response with age, we provide evidence that vascular dysfunctions in elderly subjects manifest not only as a reduction in flow magnitude, but also as temporally altered perfusion dynamics. Our simulations reinforce the notion that hypercapnia exerts a profound effect on cerebrovascular parameters, while eliciting only minor changes in CMRO_2_, supporting the assumption that hypercapnia primarily alters CBF without affecting metabolic demand. Notably, simulations showed acidic pH shifts during elevations of CO_2_, which provides a plausible link with the observation that even very mild hypercapnia (2 torr) can suppress neuronal activity in specific frequency bands (Driver et al., 2016), as recently suggested (Mangia et al., 2025 and references therein).

The alterations of vascular parameters that we report here suggest a deterioration in neurovascular coupling, a critical mechanism that ensures the matching of cerebral blood supply to neuronal activity. Impaired neurovascular coupling has been implicated in aging, with evidence pointing to early disruptions in this coupling preceding overt cognitive decline (Li et al., 2023; Tarantini et al., 2017; Zhu et al., 2022). A growing body of evidence indicates that cerebrovascular dysfunction, including altered vascular reactivity and impaired hemodynamic dynamics, can precede or accompany the earliest stages of neurodegeneration, sometimes appearing before measurable structural atrophy or canonical metabolic changes. Multimodal, large-scale data-driven analyses identify vascular dysregulation as an early abnormality in the cascade leading to late-onset dementia. For example, integrative analyses of ADNI data place vascular dysregulation ahead of amyloid, metabolic and structural markers in the temporal ordering of biomarker abnormalities (Iturria-Medina et al., 2016). Animal models likewise report endothelial dysfunction and vascular changes prior to overt amyloid plaque deposition (Waigi et al., 2024; Wang et al., 2023). Human studies show reductions in cerebrovascular reactivity (CVR) and baseline perfusion in at-risk or preclinical individuals (Chen, 2018), and indices of hemodynamic impairment (e.g., breath-holding index, CVR, baseline CBF) have been associated with subsequent cognitive decline or conversion from mild cognitive impairment to dementia (Buratti et al., 2015; Duan et al., 2021; van Dinther et al., 2024). Taken together, these findings support the proposition that alterations in temporal hemodynamic parameters (delay, time constant, gain) may reflect early vascular pathology that can antedate or contribute to later structural and metabolic deterioration towards cognitive decline.

While the reduction in CBF response amplitude (i.e., gain of the transfer function) is a known effect of aging, the changes in delay and time constant observed in older subjects reflect distinct aspects of impaired cerebrovascular dynamics. In our settings, delay captures the lag between a rise in inhaled CO_2_ and the onset of the PaCO_2_-induced CBF response, indicating the initiation of vascular adaptation. The time constant represents the rate at which the PaCO_2_-induced CBF response reaches its new steady state, reflecting the overall speed of vascular adjustment. Together, these parameters provide a complementary perspective to conventional CVR measurements, which quantify only the amplitude of flow changes (Sur et al., 2020). Our findings suggest that in aging, cerebrovascular responses are not only reduced in magnitude but also temporally affected. Results point to an increased stiffness or loss of compliant microvasculature producing a rapid but small hemodynamic change. For example, stiff vessels transmit pressure changes faster (short *τ*) but have less ability to dilate (reduced *κ*). Alternatively, or in addition, impaired endothelial signaling could reduce sustained dilation, but the immediate smooth-muscle mechanical response or pressure equilibration could be faster. Our model cannot distinguish between these alternatives with the available data, and the effect of parameter-estimation covariance cannot be excluded as in exponential-convolution models the early slope depends on the *κ*⁄*τ* ratio. We found a modest negative association between *κ* and *τ* across ROIs, and the Jacobian-derived covariance indicated appreciable coupling in the fitted parameters in several regions, consistent with a partial trade-off during optimization. However, a subset of ROIs exhibited decreased *τ* together with independent signatures of altered vascular mechanics, suggesting that both parameter-estimation covariance and genuine physiological changes may contribute. Distinguishing these mechanisms requires further targeted analyses, e.g., testing whether independent measures of vascular stiffness or CBV dynamics co-localize with decreased *τ*, which we view as important next steps.

The possible identification of altered CBF response dynamics as biomarkers for aging holds significant clinical promise. Enabling earlier detection may open opportunities for timely interventions aimed at preserving cognitive function. Moreover, monitoring these hemodynamic parameters longitudinally could facilitate the assessment of therapeutic efficacy in clinical trials targeting vascular components during the pathogenesis of cognitive decline. Incorporating dynamic hemodynamic markers into clinical workflows may enhance the precision of diagnosis and prognosis of dementia, providing a more comprehensive understanding of disease progression. However, several limitations warrant consideration besides those already mentioned. The study’s design precludes causal inferences, and the sample size may limit the generalizability of the results. Future longitudinal studies with larger cohorts are needed to validate these biomarkers and establish their prognostic value. Additionally, the integration of these dynamic hemodynamic markers with other imaging modalities, such as advanced MRI techniques (e.g., high-temporal-resolution fMRI), could provide a more holistic view of the neurovascular and metabolic alterations during aging.

In summary, we employed a biophysical model of neurovascular and neurometabolic coupling for predicting and interpreting experimental data related to hypercapnia during healthy aging. The ability to simulate complex neurovascular interactions with high fidelity represents a crucial step toward advancing our understanding of cerebral physiology and enhancing the utility of functional neuroimaging techniques.

## Acknowledgments

The authors thank Elena Russo for critical manuscripts reading; Giulia Caruso, Carlotta di Domenico, Sabrina Bonarota for participant recruitment.

## Author Contributions Statement

FG, LS, MDN conceptualized and designed research. MG, FG, TS, CM, and MC performed experiments. GG, MG, TS, EDA, MM, and MDN analyzed data. MDN developed the computational model. MDN wrote the main manuscript text. MDN prepared figures. All authors reviewed and approved the manuscript.

## Footnotes

1 Supplementary material will be available upon article publication after peer review.

2 The code repository will be released upon article publication after peer review.

3 Data requests will be considered upon article publication after peer review.

